# Insights from comparative plastid genomics of colorless facultative pathogens *Prototheca* (Chlorophyta): Unveiling membrane transport and organelle division as key functions

**DOI:** 10.1101/2024.01.16.575923

**Authors:** Kacper Maciszewski, Gabriela Wilga, Tomasz Jagielski, Zofia Bakuła, Jan Gawor, Robert Gromadka, Anna Karnkowska

## Abstract

Plastids are usually involved in photosynthesis, but the secondary loss of this function is a widespread phenomenon in various lineages of algae and plants. In addition to the loss of genes associated with photosynthesis, the plastid genomes of colorless algae are frequently reduced further. To understand the pathways of reductive evolution associated with the loss of photosynthesis, it is necessary to study a number of closely related strains. *Prototheca*, a chlorophytean genus of facultative pathogens, provides an excellent opportunity to study this process with its well-sampled array of diverse colorless strains.

We have sequenced the plastid genomes of 13 *Prototheca* strains and reconstructed a comprehensive phylogeny that reveals evolutionary patterns within the genus and among its closest relatives. Our robust phylogenomic analysis revealed three independent losses of photosynthesis among the *Prototheca* strains and considerable coding content variability in their ptDNA. Despite this diversity, all *Prototheca* strains retain the same key plastid functions. These include processes related to gene expression, as well as crucial roles in fatty acid and cysteine biosynthesis, membrane transport, and organelle division. While the retention of vestigial genomes in colorless plastids is typically associated with the biosynthesis of secondary metabolites, the remarkable conservation of plastid membrane transport and organellar division systems in the nonphotosynthetic genera *Prototheca* and *Helicosporidium* provides an additional constraint against the loss of ptDNA in this lineage. Furthermore, these genes can potentially serve as targets for therapeutic intervention, indicating their importance beyond the evolutionary context.

## 1. Introduction

Plastids are eukaryotic organelles derived from cyanobacteria, whose most widely recognized and deeply studied function is photosynthesis (Archibald, 2009; de Vries and Archibald, 2017). However, even though carrying the photosynthetic apparatus is their distinctive feature, a variety of biochemical pathways have been inherited by the extant plastids from their cyanobacterial ancestor, and their functions, such as heme, fatty acid or amino acid biosynthesis, remain crucial constituents of the hosts’ metabolism (Lim and McFadden, 2010; Maciszewski and Karnkowska, 2019; Novák Vanclová et al., 2020). As a result, the rather common occurrence of photosynthesis loss does not necessarily lead to the disappearance of the organelle – on the contrary, the so-called colorless plastids can be found in almost all known plastid-bearing lineages (Hadariová et al., 2018; Maciszewski and Karnkowska, 2019), with only a handful of known cases of photosynthesis loss leading to a radically different outcome (Gornik et al., 2015; Janouškovec et al., 2017).

With certain exceptions, even the most functionally reduced plastids retain vestigial forms of genomes (ptDNA), whose coding contents are usually limited to up to several metabolically relevant protein-coding genes and a minimal transcription and translation apparatus (Figueroa-Martinez et al., 2014; Kayama et al., 2020; Smith, 2018a; Smith, 2018b; Smith and Lee, 2014). Despite their small size and modest genetic repertoire, the genomes of colorless plastids can be important sources of insight into the evolutionary history and lifestyle of their host, as well as, in case of plastid-bearing parasites, even their vulnerabilities (Kayama et al., 2020; Lim and McFadden, 2010; Salomaki and Kolísko, 2019; Sibbald and Archibald, 2020).

To understand the plastid (and plastome) evolution in secondarily non-photosynthetic organisms, comparative genomic and transcriptomic analyses with their photosynthetic relatives are the most common *modus operandi*. This applies both to lineages where photosynthesis losses have repeatedly occurred late in their evolution, such as orchids (Barrett et al., 2019), dinoflagellates (Hoon Kim et al., 2013; Janouškovec et al., 2017) or the diatom genus *Nitzschia* (Kamikawa et al., 2015b), and those whose shift toward heterotrophy preceded their major radiation, such as apicomplexans (Salomaki and Kolísko, 2019; Sato, 2011). Close relatedness between organisms exhibiting vastly different lifestyles is often a hallmark of complex and captivating evolutionary paths, and a perfect example of that can be found among the green algal order Chlorellales. The relatives of the model green microalga *Chlorella* include a photosynthetic genus *Auxenochlorella*, as well as two secondarily non-photosynthetic genera – *Helicosporidium*, which are highly specialized, obligate gut parasites of insects (de Koning and Keeling, 2006; De Koning and Keeling, 2004; Sun and Pombert, 2014; Tartar, 2013), and *Prototheca*, which are predominantly free-living opportunistic pathogens of diverse vertebrates, including humans (Borza et al., 2005; Guo et al., 2022; Jagielski et al., 2019, 2018). The evolutionary past of these organisms, especially their transitions between photosynthetic, parasitic and free-living heterotrophic lifestyles, remains mysterious even with the availability of several genomic datasets.

The aforementioned assemblage, collectively referred to as the AHP clade (Suzuki et al., 2018), constitutes a rather unique model for studying evolutionary transitions related to plastid reduction. This is because non-photosynthetic primary plastids often tend to adopt extreme forms, as demonstrated by cases of ptDNA inflation in certain green algae, such as *Leontynka pallida* or *Polytoma uvella* (Figueroa-Martinez et al., 2016; Pánek et al., 2022), as well as complete ptDNA loss in other chlorophytes (*Polytomella parva*) and even land plants (*Rafflesia* sp.) (Molina et al., 2014; Smith and Lee, 2014). On the other hand, genomes of colorless plastids that are reduced in size and function, while retaining a rudimentary set of genes associated with metabolite synthesis and housekeeping functions, are more typical for the substantially better-studied secondary plastids, found e.g. in diatoms or apicomplexans (Kamikawa et al., 2018; Lim and McFadden, 2010). What is more, *Prototheca* and *Helicosporidium* are also among the extremely rare primary plastid-bearing pathogens, which makes the convergently similar form of their ptDNA to apicoplast genomes even more interesting. However, although *Prototheca* infections have been repeatedly observed in humans, dogs, and cows, its occurrence in a vast variety of other vertebrate hosts has been documented almost entirely in single case studies (Jagielski et al., 2019); the transmission, infectivity, and mechanisms of pathogen-host interactions therefore remain unknown (Shave et al., 2021). As demonstrated by the past studies of Apicomplexa, understanding the functions of vestigial plastids in parasites can not only provide key insights into their metabolic dependence on the host (Sheiner et al., 2013), but also uncover potential targets for therapeutical agents (Mukherjee and Sadhukhan, 2016).

## 2. Materials and methods

### 2.1 Cultivation, DNA isolation and sequencing

18 strains of *Prototheca* cultured on agar plates have been obtained from the in-house culture collection of the Department of Medical Microbiology (Institute of Microbiology, Faculty of Biology, University of Warsaw, Poland). Their DNA isolation was performed according to the optimized protocol outlined in (Jagielski et al., 2017), and the samples were sequenced using Illumina MiSeq PE300 platform, with 600-cycle chemistry kit.

### 2.2 Quality control and genome assembly

Quality control of the obtained reads was carried out using FastQC v0.11.5 (Andrews, 2010). Adapter removal and trimming was performed using Trimmomatic v0.32 (Bolger et al., 2014) using default parameters. Initial assembly was carried out using SPAdes v3.11.1 (Bankevich et al., 2012), and the outputs were analyzed to assess their general quality using Quast v5.0.2 (Gurevich et al., 2013). The detection of potential contamination was done using Blobtools v1.1 (Laetsch and Blaxter, 2017).

Among the assembled contigs, plastid genome-derived sequences were identified using Tiara v1.01 (Karlicki et al., 2022), supplemented by BLASTn searches (Altschul et al., 1990) using publicly available ptDNA sequences of *Prototheca* spp. as queries. Largest identified plastid genome fragments in each assembly were extracted and subsequently used as seeds for final ptDNA assembly using NOVOPlasty v4.3 (Dierckxsens et al., 2017). Circularized plastid genomes were recovered in all 18 datasets; five of them were identical with already published plastid genome sequences.

### 2.3 Plastid genome annotation and visualization

Automatic annotation of *Prototheca* plastid genomes was carried out using Geneious Prime v2022.1.1 software (https://www.geneious.com) using Live Annotate & Predict toolkit (Find ORFs and Annotate From… features), utilizing a manually constructed database of published plastid genomes of *Prototheca* spp., *Chlorella* spp., *Auxenochlorella* spp. and *Parachlorella kessleri*. Identities of all protein-coding gene sequences were confirmed by alignment with the NCBI non-redundant protein database (NCBI-nr) via BLASTX algorithm (Altschul et al., 1990) and with the PFAM 35.0 protein families’ database (pfam.xfam.org) using the browser-accessible internal HMM search feature (Mistry et al., 2021). Plastid genome maps were generated using the OGDraw v1.3.1 online tool (Greiner et al., 2019).

### 2.4 Plastid genome-based phylogenomic analysis

79 protein-coding genes were extracted from the 40 annotated ptDNA sequences of *Prototheca* and their closest relatives, including: the 13 *Prototheca* strains analyzed in this work (five of the newly obtained genomes were identical with previously sequenced ones), six other published plastid genomes of *Prototheca*, 11 published plastid genomes of *Chlorella*, two published plastid genomes of *Auxenochlorella*, two published plastid genomes of *Pedinomonas*, and the single published plastid genomes of *Helicosporidium* sp., *Marsupiomonas* sp., *Dicloster acuatus, Marvania geminata, Parachlorella kessleri*, and *Pseudochloris wilhelmii*. All coding sequences were translated into amino acid sequences, aligned using L-INS-I method in MAFFT v7.310 (Katoh and Standley, 2013), and concatenated using catsequences script (https://github.com/ChrisCreevey/catsequences) to produce a data matrix with total length of 33,713 amino acids.

The concatenated alignment was used as the input for phylogenetic analyses via maximum likelihood method implemented in IQ-TREE v2.0.6 software (Minh et al., 2020), and via Bayesian inference method implemented in MrBayes v3.2.6 (Ronquist et al., 2012). Maximum likelihood phylogeny reconstruction used a partitioned matrix with automatic substitution model selection for each partition (*-m TEST* parameter), and 1000 non-parametric bootstrap replicates. Bayesian reconstruction used a non-partitioned dataset with preset sequence evolution model (cpREV), with 1 000 000 generations (incl. 250 000 generations burn-in), after which convergence of the Markov chains was achieved. Both methods yielded mostly congruent tree topology, with local divergence in topologies described in further detail in the Results and discussion section.

### 2.5 Rate of evolution estimation

Codon alignments of 25 plastid protein-coding genes were prepared using PAL2NAL v14 software (Suyama et al., 2006). Rates of synonymous and non-synonymous substitutions (*dN/dS*) for all gene alignments were calculated using CodeML tool implemented in PamlX v1.3.1 toolkit (Xu and Yang, 2013). Mean *dN/dS* values were calculated for two groups: *Prototheca* clade B (7 taxa) and clade C (11 taxa) for all 25 protein-coding genes identified in the ptDNA sequences obtained for these taxa, and compared using two-sided Mann-Whitney *U*-test implemented in Social Science Statistics calculator (online tool; https://www.socscistatistics.com/tests/mannwhitney/).

## 3 Results and discussion

### 3.1 Plastid-based phylogeny of the genus *Prototheca*

Plastid genome characteristics of *Prototheca* spp. are shown in Table 1. The plastid genome-based phylogenetic tree of *Prototheca* spp. and their relatives is shown on Figure 1. Despite the overall high support for the reconstructed phylogeny, both estimated by Bayesian posterior probability and bootstrap support values, we observed two topological incongruencies between the Bayesian and maximum likelihood reconstructions. First, the branching order of *Prototheca fontanea* and *P. lentecrescens* in the ML reconstruction is reversed in comparison with the Bayesian tree, which resolved the topology as *P. lentecrescens* branching off earlier than *P. fontanea* (see the dotted line-framed section of the clade B on Figure 1). Secondly, *P. moriformis* branches off at the base of the *P. tumulicola, P. blaschkeae* and *P. stagnora* clade in the ML reconstruction, instead of the base of its sister clade encompassing *P. bovis, P. cerasi, P. ciferrii, P. zopfii, P. pringsheimii, P. vistulensis* and *P. cookei* (see the dotted line-framed section of the clade C in Figure 1). In both cases, however, the Bayesian topology is further supported by the gene content of the plastid genomes of the *Prototheca* strains in question – as a single loss of a gene set (the six-gene *atp* family in case of the clade B, and the four-gene *rpo* family in case of the clade C; Figure 1) is logically a more likely occurrence than two independent losses of the same gene set in closely-related lineages, therefore we present the Bayesian reconstruction on the Figure 1 as the more evolutionary plausible.

**Table 1.** Plastid genome characteristics of *Prototheca* spp. Taxa whose plastid genomes were first sequenced in this study are distinguished in **bold**.

| Organism | ptDNA size<br>(bp) | Protein-coding<br>genes | Accession<br>no. |
| --- | --- | --- | --- |
| *Prototheca xanthoriae* SAG263-11 | 55,636 | 40 | KJ001761 |
| *Prototheca cutis* ATCC PRA-338 | 51,673 | 40 | NC037480 |
| *Prototheca wickerhamii* DBVPG | 47,997 | 35 | NC054192 |
| *Prototheca stagnora* ATCC16528 | 48,253 | 29 | NC037479 |
| *Prototheca bovis* SAG2021 | 28,638 | 19 | MF197536 |
| *Prototheca ciferrii* SAG2063 | 28,698 | 19 | MF197535 |
| ***Prototheca paracutis* YMTW3-1** | 51,694 | 40 |  |
| ***Prototheca miyajii* IFM53848** | 53,237 | 40 |  |
| ***Prototheca lentecrescens* PK1** | 53,163 | 40 |  |
| ***Prototheca fontanea* PK2** | 46,196 | 34 |  |
| ***Prototheca wickerhamii* PK9** | 47,600 | 35 |  |
| ***Prototheca tumulicola* JCM31123** | 49,137 | 26 |  |
| ***Prototheca blaschkeae* SAG2064** | 46,294 | 30 |  |
| ***Prototheca zopfii* ATCC16533** | 28,349 | 19 |  |
| ***Prototheca cerasi* JCM9400** | 28,412 | 19 |  |
| ***Prototheca pringsheimii* SAG263-3** | 28,370 | 19 |  |
| ***Prototheca vistulensis* W3** | 28,516 | 19 |  |
| ***Prototheca cookei* ATCC16527** | 28,282 | 19 |  |
| ***Prototheca moriformis* SAG263-2** | 38,525 | 22 |  |

**Figure 1.**
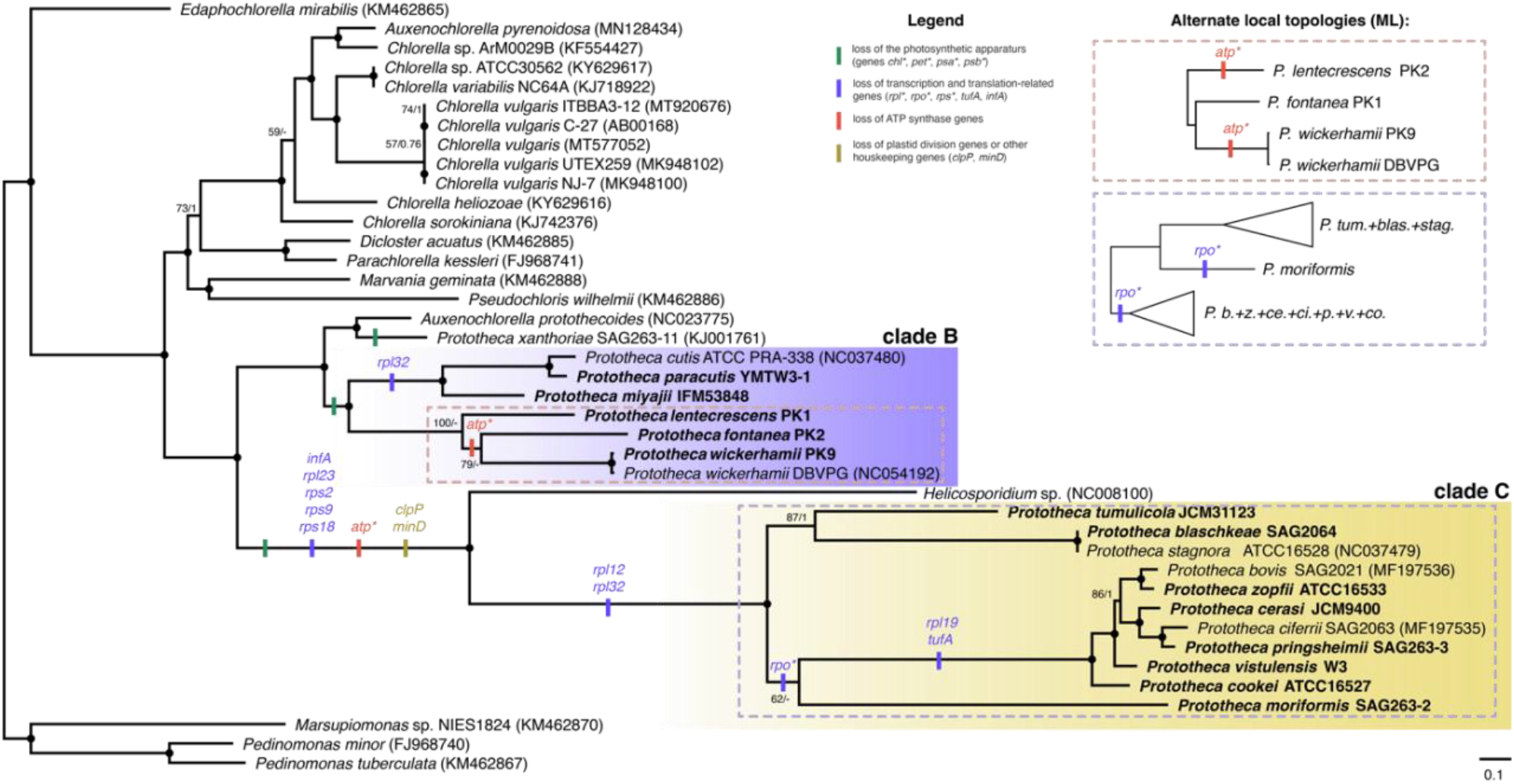
Phylogenomic analysis of the *Prototheca* and other Chlorellales with mapped gene losses on the respective branches. Tree shown is a Bayesian inference (BI) phylogeny of a 79 marker genes from plastid genomes. Maximum likelihood (ML) phylogeny was congruent with BI with two exceptions shown. The phylogeny is based on a concatenated marker gene alignment of 33,713 unambiguously aligned sites under the model cpREV+G4. Black dots indicate maximal support for a particular node. When not maximal, only *a posteriori* >0.5 and bootstrap support values >50% are shown. Strain names in **bold** denote ptDNA sequences obtained in this study. NCBI GenBank accession numbers for publicly available sequences are provided in brackets.

Regardless of the incongruencies described above, both methods resolve the three main *Prototheca* clades identically, with the first clade (represented solely by *P. xanthoriae*, formerly classified as *P. wickerhamii* (Borza et al., 2005)) branching off as sister to *Auxenochlorella protothecoides*, the second clade (encompassing *P. cutis, P. paracutis, P. miyajii, P. wickerhamii, P. fontanea* and *P. lentecrescens*; further referred to as “*Prototheca* clade B”) branching as sister to the *A. protothecoides* + *P. xanthoriae* clade, and the third (encompassing all remaining species: *P. tumulicola, P. blaschkeae, P. stagnora, P. moriformis, P. bovis, P. zopfii, P. cerasi, P. ciferrii, P. pringsheimii, P. vistulensis* and *P. cookei;* further referred to as “*Prototheca* clade C”) branching off as sister to *Helicosporidium* sp. As the last common ancestor of all *Prototheca* spp. is also the ancestor of *Helicosporidium* sp. and *Auxenochlorella protothecoides*, the genus *Prototheca* is therefore, by definition, polyphyletic, as suggested in the previous nuclear and mitochondrial gene-based phylogenies (Jagielski et al., 2019, 2018). Furthermore, it is clear from the plastid genome-based phylogeny presented above that all three *Prototheca* clades originate from ancestors which lost their photosynthetic capabilities independently.

However, due to vast discrepancies in sampling across the diversity of *Prototheca* spp. between the aforementioned phylogenies and the current work, we believe them to be impossible to compare in detail – for instance, the most recent *cytb*-based reconstruction included seventeen *P. ciferrii* sequences, while the species *P. paracutis* or *P. lentecrescens* have not been included at all, simply because they have not been described at the time (Jagielski et al., 2022, 2019; Kunthiphun et al., 2019).

### 3.2 Independent losses of the photosynthetic apparatus shaped divergent paths of *Prototheca* plastid genomes

Interestingly, it is noticeable that the plastid genome contents differ substantially between the three *Prototheca* clades. While the ptDNA of *P. xanthoriae* is both largest (over 55.6 kbp) and *ex aequo* most gene-rich (40 protein-coding genes) of all *Prototheca* spp., the size and gene contents across the closely-related *Prototheca* clade B are generally comparable, ranging from approximately 48.0 kbp and 35 protein-coding genes in *P. wickerhamii* to approximately 53.2 kbp and 40 protein-coding genes in *P. lentecrescens*. In contrast, the plastid genomes of the *Prototheca* clade C carry no more than 26 protein-coding genes, with many species (e.g. *P. zopfii, P. bovis*) having only 19. Moreover, the diminished gene content of the ptDNA in *Prototheca* clade C is not entirely proportional to their length –the plastid genomes of *P. tumulicola* and *P. stagnora* exceed 48.0 kbp in size, which would be within the range of *Prototheca* clade B, but their substantially smaller gene content indicates elevated proportion of the non-coding regions in their plastid genomes. What is more, the ptDNA of *P. cookei* is below 28.3 kbp in size, which makes it, along with its closest relatives – *P. zopfii, P. ciferrii, P. bovis, P. pringsheimii, P. vistulensis* and *P. cerasi*, all with ptDNA length below 29 kbp – the carrier of possibly the most reduced plastid genome among unicellular eukaryotes, with only a few species of mycoheterotrophic and parasitic plants reaching smaller ptDNA size and gene contents (Smith, 2018).

Furthermore, the reduction of coding contents of *Prototheca* plastomes is definitely not a symptom of their random decay in time, but a manifestation of divergence of function, as indicated by the retention or losses of complete gene operons or families, such as *atp* and *rpo*. Although the documentation for loss of these two families in non-photosynthetic plastids is rather abundant, pointing toward the possibility of functional compensation for the missing *rpo* by nuclear RNA polymerases (Börner et al., 2015; Chen et al., 2020; Graham et al., 2017; Mohanta et al., 2020), the retention of ATP synthase subunits in *P. xanthoriae, P. cutis, P. paracutis, P. miyajii* and *P. lentecrescens* indicates their capability for generating ATP or alternatively, proton motor force across the plastid membrane. Although the ATP synthase subunits have been identified among the ptDNA contents of non-photosynthetic cryptophytes (Donaher et al., 2009), diatoms (Kamikawa et al., 2015a), and even land plants (Logacheva et al., 2014), their role in absence of photosynthesis still awaits full explanation. Still, the presence of *atp* genes suggests that the plastids of the aforementioned five representatives of *Prototheca* carry out certain currently inscrutable metabolic processes absent from all the others. It might additionally be worth noting that all six plastid-encoded *atp* genes exhibit significantly increased rate of evolution (expressed as *dN/dS* values) in *Prototheca* compared to the photosynthetic Chlorellales (Mann-Whitney *U-*test; *p*-values between 1.83·10^−5^ and 0.028).

Bearing this functional diversity in mind, it is tempting to hypothesize whether the independent losses of photosynthesis in the ancestors of the three *Prototheca* clades might have been the cornerstone behind their divergence. A factor that certainly has to be taken into account is time – with the plastid genome of *P. xanthoriae* lacking only the photosynthetic apparatus (and therefore displaying rather “basic” reduction), compared to the phototrophic Chlorellales, and the ptDNA of many *Prototheca* clade C representatives being reduced to just a handful of genes (and therefore displaying “advanced” reduction), one could assume that the loss of photosynthesis in the clade C’s ancestor occurred earlier than in those of clade B and *P. xanthoriae*. Such hypothesis, however, might be quite difficult to prove, as calibration of the evolutionary timeline would require insight into the fossil record. This, on the other hand, would be rather challenging not only due to the scarcity of adequately conserved fossilized remains of non-skeleton-forming unicellular eukaryotes, but also because of the near-identical morphology of all extant *Prototheca* species, which would make the phylogenetic placement of an extinct one nearly impossible.

### 3.3 Membrane transport and regulation of organelle division are the previously overlooked constraints against genome loss in colorless plastids

Despite the differences outlined above, plastid genomes of *Prototheca* spp. (and *Helicosporidium* sp.) also share a vast array of similarities – all examined species retained a common core set of 19 genes, covering the entire plastid gene repertoire of certain clade C representatives, such as the aforementioned *P. cookei*. This includes 15 genes involved in transcription and translation, but also a chloroplast division-associated gene *ftsH*, a fatty acid synthesis-associated gene *accD*, and two membrane transport machinery components: *ycf1*, encoding the largest subunit of the protein translocation system TIC (TIC214), and *cysT*, involved in sulfate ion import across the plastid membrane (Figure 2).

**Figure 2.**
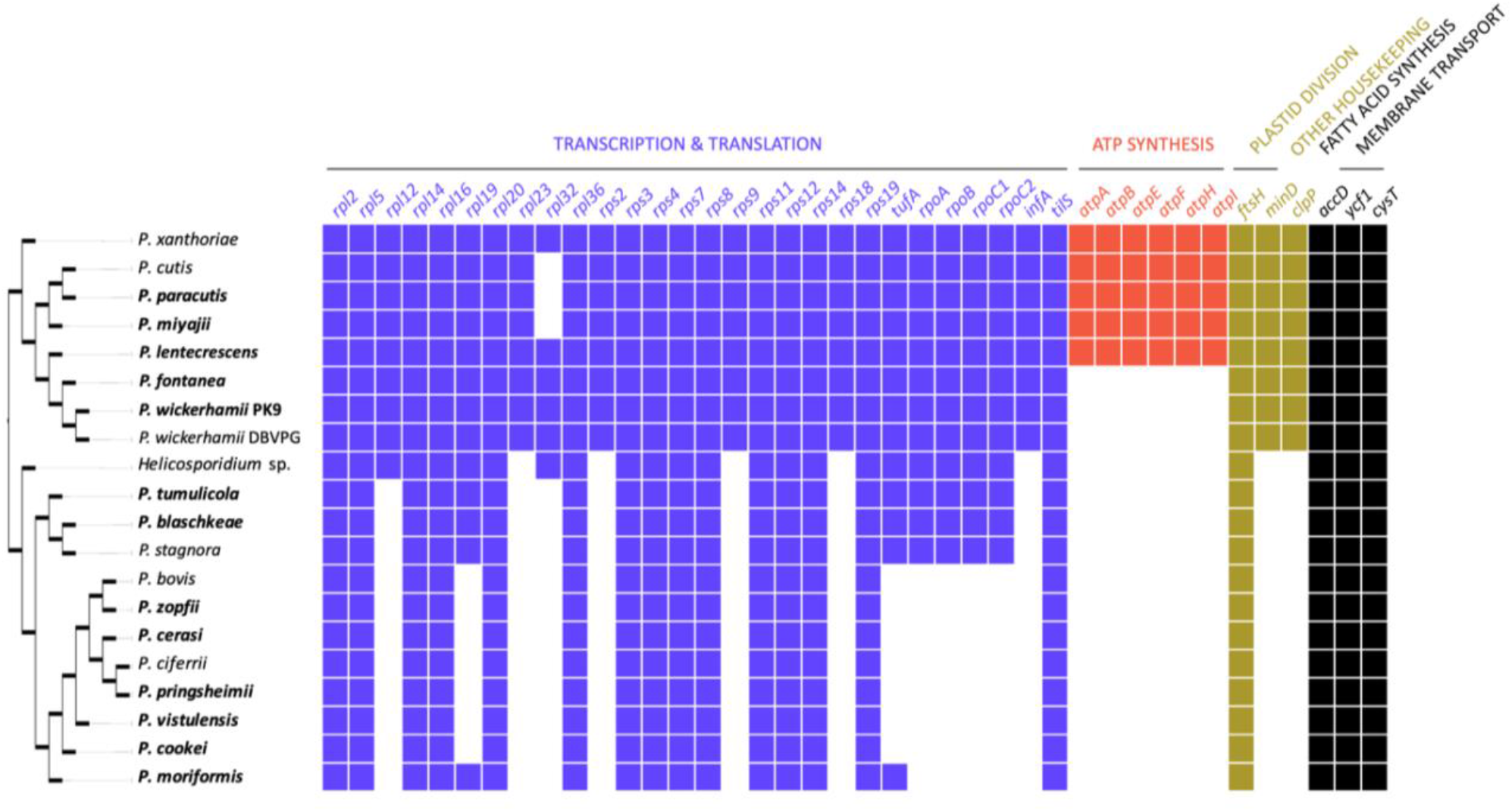
Distribution of various plastid genome functions across *Prototheca* spp.

The available body of evidence suggests that the driving force behind genome retention in non-photosynthetic plastids is almost invariably the presence of indispensable plastid-encoded secondary metabolite synthesis pathway components (Kayama et al., 2020; Maciszewski and Karnkowska, 2019). In contrast, the role of *ftsH* and *ycf1* is evidently the maintenance of transport and division mechanisms of a plastid compartment that serves almost entirely nucleus-encoded biosynthetic pathways, such as amino acid, heme or fatty acid synthesis, demonstrated in past studies to be carried out in the vestigial plastids of *Helicosporidium* strain AT-2000 (De Koning and Keeling, 2004) and *Prototheca xanthoriae* strain SAG263-11 (Borza et al., 2005), and now found to have only two plastid-encoded components (*accD* and *cysT*) in total.

This work is not the first report of *ycf1* and *ftsH* retention in the genomes of non-photosynthetic plastids; both were reported in the previously studied plastid genomes of *Helicosporidium* and certain *Prototheca* spp., as well as the distantly related non-photosynthetic, although free-living chlorophyte *Polytoma uvella* (de Koning and Keeling, 2006; Figueroa-Martinez et al., 2016; Severgnini et al., 2018; Suzuki et al., 2018). However, the potentially key role of *ycf1* may have been overlooked in the past, as the first reports of its retention in non-photosynthetic plastids (de Koning and Keeling, 2006) predate the discovery of this gene’s biological role (de Vries et al., 2015; Nakai, 2015) – hence its name still suggests it to be a gene of unknown function. What is more, *ycf1* is a plastid-encoded gene exclusively in land plants and chlorophytes, while the bulk of studies on roles of non-photosynthetic plastids is focused on secondary plastid-bearing lineages, such as apicomplexans (Hadariová et al., 2018).

Still, the plastid genome contents that are not shared by all *Prototheca* spp. remain quite mysterious, especially the ATP synthase operon. Although its inconsistent presence in this genus has been pointed out before (Suzuki et al., 2018), the broader sampling of our study made it possible to observe that the entire *atp* gene set was differentially lost among *Prototheca* clade B, which has also been documented to occur in certain land plants, such as Orobanchaceae (Chen et al., 2020; Wicke et al., 2013), but in contrast with its consistent retention or loss in the descendants of all photosynthesis loss events in secondary plastids (Kamikawa et al., 2015a; Suzuki et al., 2018). In non-photosynthetic plastids, the proposed role of the ATP synthase complex is the hydrolysis of ATP to generate proton motive force across the inner plastid membrane, which is utilized for protein translocation by the twin-arginine translocase (Tat) system (Kamikawa et al., 2015a). However, while the Tat system subunits have been identified in other non-photosynthetic lineages that retain ptDNA-encoded ATP synthase complex, e.g. *Leontynka pallida* (Chlorophyta) or *Cryptomonas paramecium* (Cryptophyta), it is absent both in plastid and nuclear genomes of all *Prototheca* investigated so far (Donaher et al., 2009; Pánek et al., 2022; Suzuki et al., 2018). Therefore, the ATP synthase presence in some of the *Prototheca* spp. could be explained by the existence of an unknown protein translocation system that relies on the proton gradient, or a completely different, *Prototheca*-specific role of this complex in plastids, as proposed by Suzuki et al. 2018. Nonetheless, it is almost certain that there is a variability in plastid functions among *Prototheca* that cannot be fully explained by their plastome-encoded components.

Furthermore, the analysis of the rates of evolution of ptDNA-encoded genes between *Prototheca* clades yielded rather unexpected results (Table 2). Among 25 analysed genes, 15 (*accD, ftsH, rpl12/14, rpoA/B/C1, rps2/3/4/8/14/18, tufA, ycf1*) displayed significantly increased *dN/dS* values in *Prototheca* clade C, compared to clade B; eight others (*rpl5/16/32/36, rpoC2, rps11/12, tilS*) exhibited no difference between clades, but most surprisingly, two genes – *cysT* and *rpl20* – have apparently undergone accelerated evolution in the *Prototheca* clade B, compared to clade C. This might be indicative of diversified evolutionary pressure toward different gene (and protein) sequence conservation between the *Prototheca* clades, with e.g. *accD* and *ycf1* undergoing more constrained evolution in the *Prototheca* clade B, and *cysT* being more conserved in the clade C.

**Table 2.**
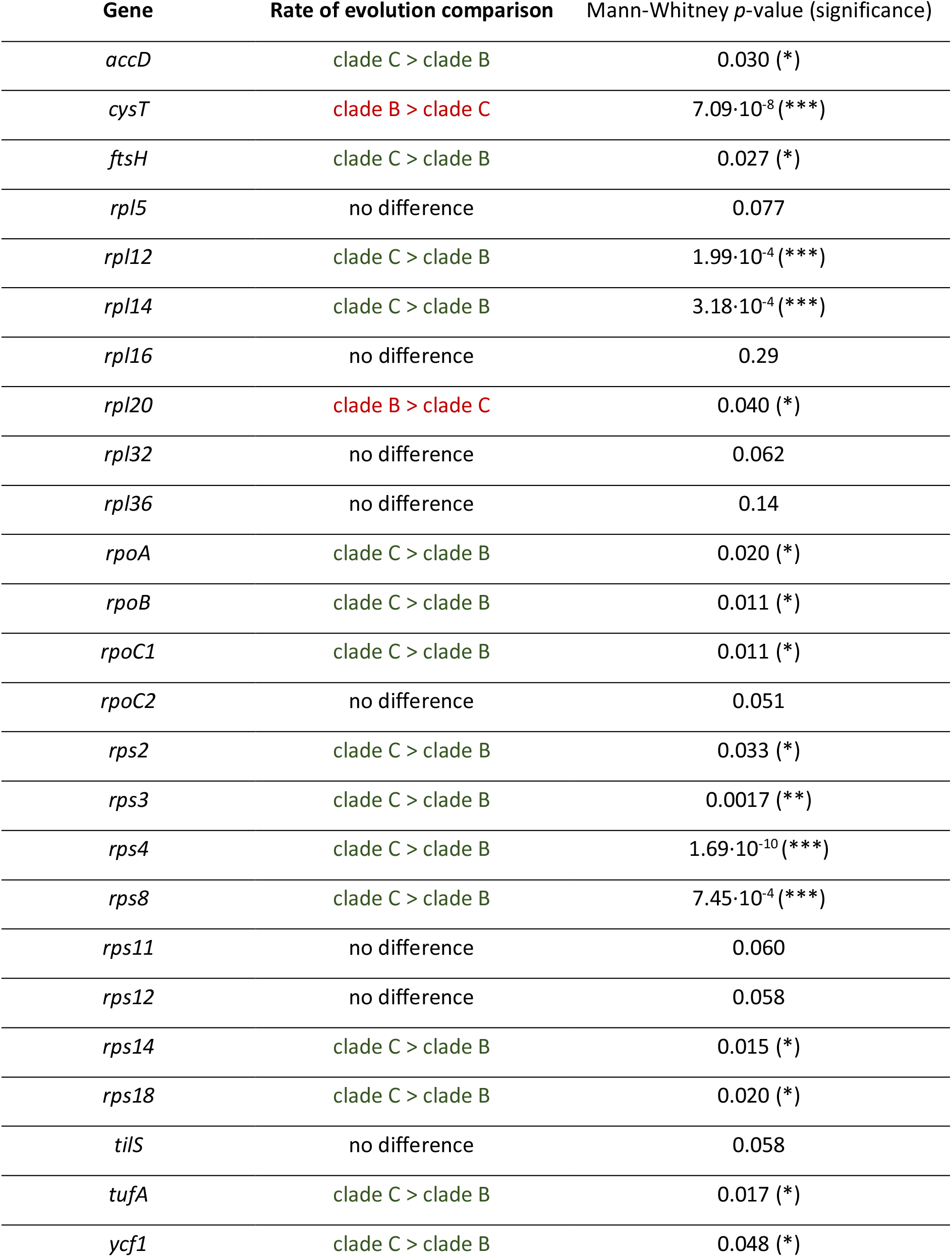
Comparative analysis of the rate of evolution (*dN/dS*) of protein-coding genes in the plastid genomes of two main *Prototheca* clades.

Interestingly, despite the observed accelerated evolutionary rate and gene losses in the ptDNA of different *Prototheca* lineages, we have not identified a single symptom of pseudogenization, i.e. disruption of a reading frame in a discernible protein-coding gene. This stands in contrast with a wide array of past studies (Barrett et al., 2014; Bellot and Renner, 2016; Wicke et al., 2013), in which pseudogenes have been frequently identified in non-photosynthetic primary plastid genomes, especially those of land plants, and have been considered the hallmark intermediate stages in the gradual reductive evolution of ptDNA.

## 4 Conclusions

In this study, we obtained 13 new complete plastid genome sequences of *Prototheca* spp. – a paraphyletic assemblage of secondarily non-photosynthetic representatives of Chlorellales. We have demonstrated that despite having forfeited the photosynthetic apparatus three times independently and bearing highly variable coding contents, the plastid genomes of all *Prototheca* share the same key functions, which, apart from own gene expression-related processes, include fatty acid and cysteine biosynthesis, as well as membrane transport and organelle division. Although components of the pathways involved in secondary metabolite biosynthesis have been demonstrated to be the crucial factor behind the retention of vestigial genomes in colorless plastids of many eukaryotic lineages, the plastid membrane transport and organellar division systems being additional constraint against ptDNA loss in *Prototheca* and *Helicosporidium* makes these peculiar chlorophytes a prominent exception from the general paradigm. What is more, the retention of genes unique to plastids in these opportunistic pathogens could be exploited in a clinical setting, as therapeutical agents targeting plastid transport machinery, such as the product of the gene *ycf1*, would likely pose minimal risk for the patients.

Nonetheless, we are convinced that the facultatively pathogenic inclinations of *Prototheca* have not been the main driving force between the repeated loss of photosynthesis. Instead, it seems more plausible that forfeiting photosynthesis accompanied the transition of these microorganisms to low-light, high-TOC habitats, such as river sediments, demonstrated in a recent environmental survey (Jagielski et al., 2022) to be their important reservoir in nature. Still, as the *Prototheca* spp. seem to possess no obvious novel adaptations both to the benthic and the pathogenic lifestyle, it would be reasonable to perceive these organisms as ecological opportunists, losing excessive genetic and biochemical burdens over the course of evolution to limit the energetic expenses of survival.

Further genomic and transcriptomic studies are necessary to explain the diversity of ptDNA contents in *Prototheca* spp. and its possible correlation with diversified plastid metabolism. We believe that the key to unraveling the mystery behind the divergence among *Prototheca* may be understanding the role of the retained ATP synthase in these organisms’ plastids, and that the *Auxenochlorella/Helicosporidium/Prototheca* assemblage could become a promising model for future studies on divergent evolution of the endosymbiotic organelles, including, but not limited to the primary plastids.

## 5 Acknowledgements

This work was supported by the European Molecular Biology Organization and Polish Ministry of Education and Ministry of Education and Science, Poland [EMBO Installation Grant 4150 to AK], National Science Centre, Poland [Sonata grant 2016/21/D/NZ8/01288 to AK], and National Science Centre, Poland [PRELUDIUM grant 2013/09/N/NZ2/00248 to ZB].

## 6 Author contributions

AK and TJ conceived and planned the project. ZB cultured all *Prototheca* strains, JG and RG isolated DNA and performed sequencing experiments including initial data curation. KM carried out bioinformatic and phylogenetic analyses, and wrote the first draft of the manuscript. GW participated in the assembly and annotation of data. KM and AK wrote the final version of the manuscript with input of other co-authors. All authors read, critically commented and approved the final manuscript.

